## Supplementary Figures S1-S12 for "Membrane curvature governs the distribution of Piezo1 in live cells"

**The supplement contains:**

Materials

Supplementary Figures S1-S12

**Materials**

Chemicals

| **Name** | **Source** | **Catalog Number** |
| --- | --- | --- |
| EMEM | Fisher scientific | MT10009CV(Corning™) |
| DMEM | Fisher scientific | 11965092  11960051 |
| FBS | Fisher scientific | FB12999102(Fisherbrand™)  FB-12 (Omega Scientific) |
| Sodium Pyruvate | Fisher scientific | 11360070 |
| Penicillin-Streptomycin | ThermoFisher | 15070063(Gibco™)  15140122(Gibco™) |
| PBS | ThermoFisher | 10010023(Gibco™) |
| Trypsin | ThermoFisher | 25300054(Gibco™) |
| TrypLE Express | ThermoFisher | 12604013 |
| Opti-MEM | ThermoFisher | 31985070(Gibco™) |
| TransIT-X2 | Mirus | MIR6003(Mirus) |
| Dulbecco's PBS | Gibco | 14-190-250 |
| GlutaMax | ThermoFisher | 35050-061 |
| sodium pyruvate | ThermoFisher | 11360-070 |
| non-essential amino acid solution | ThermoFisher | 11140-050 |
| Lipofectamine 3000 | ThermoFisher | L3000001 |
| NaCl | Fisher scientific | BP358-212(Fisher BioReagents) |
| KCl | Millipore Sigma | 7300-500GM(Millipore) |
| HEPES | Fisher scientific | BP310-1(Fisher BioReagents) |
| Glucose | Fisher scientific Sigma-Aldrich | BP350-1(Fisher BioReagents)  G-6152 |
| CaCl_2_ | Fisher scientific | C79-500 |
| MgCl_2_ | Fisher scientific | M33-500 |
| Yoda1 | Millipore Sigma | SML1558-5MG(Sigma-Aldrich) |
| Latrunculin B | AdipoGen | AG-CN2-0031-M001(AdipoGen) |
| 4.5 μm bead | Spherotech | DIGP-40-2 |
| Matrigel | Fisher scientific | 354234 |
| Gelatin | Fisher scientific | ES-006-B |
| PDL | Millipore Sigma | P7405-5MG(Sigma-Aldrich) |
| DMSO | Fisher scientific | BP231-100 |

Plasmids

| **Name** | **Source** | **Catalog #** |
| --- | --- | --- |
| eGFP-CaaX | Adam Cohen lab | N.A. |
| GPI-eGFP | Addgene | 32601 |
| GPI-mCherry | Addgene | 127812 |
| hPiezo1-eGFP | Charles Cox lab, eGFP fused at position 1591 of human Piezo1 | N.A. |
| hPiezo1-mCherry | Charles Cox lab, mCherry fused at position 1591 of human Piezo1 | N.A. |
| mOrange2-CaaX | Adam Cohen lab | N.A. |
| mPiezo1-mCherry | Ardem Patapoutian lab, mCherry fused at the C-terminus of mouse Piezo1 | N.A. |
| mTREK1-mCherry | Philip Gottlieb lab | N.A. |
| SNAP-D2R-eGFP | Adam Cohen lab | N.A. |

**Figure S1. hPiezo1-eGFP is localized to the plasma membrane**

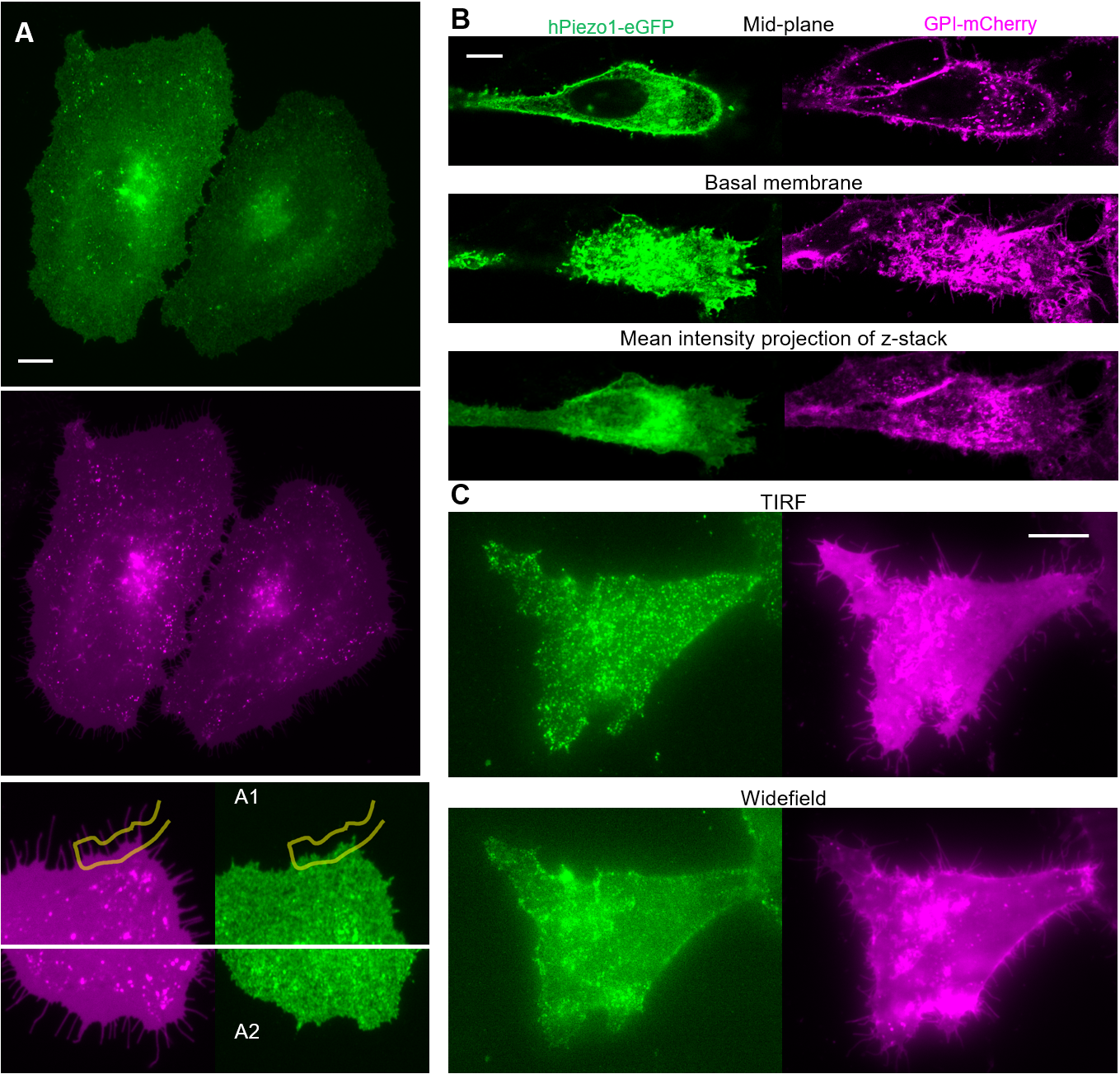

Fluorescence images of HeLa cells co-expressing hPiezo1-eGFP (green) and GPI-mCherry (magenta). **(A)** Widefield fluorescent images in Fig. 1A under normal contrast. Zoom-in images for regions A1 and A2 (marked in Fig, 1A) are shown below. **(B)** Confocal images at the middle (up) and bottom planes (middle) of a cell. Mean intensity projection of the cell shown at the bottom. Imaging was performed on a Zeiss Axio Observer 7 confocal microscope. **(C)** TIRF (up) and corresponding widefield images of the same cell. All scale bars are 10 µm.

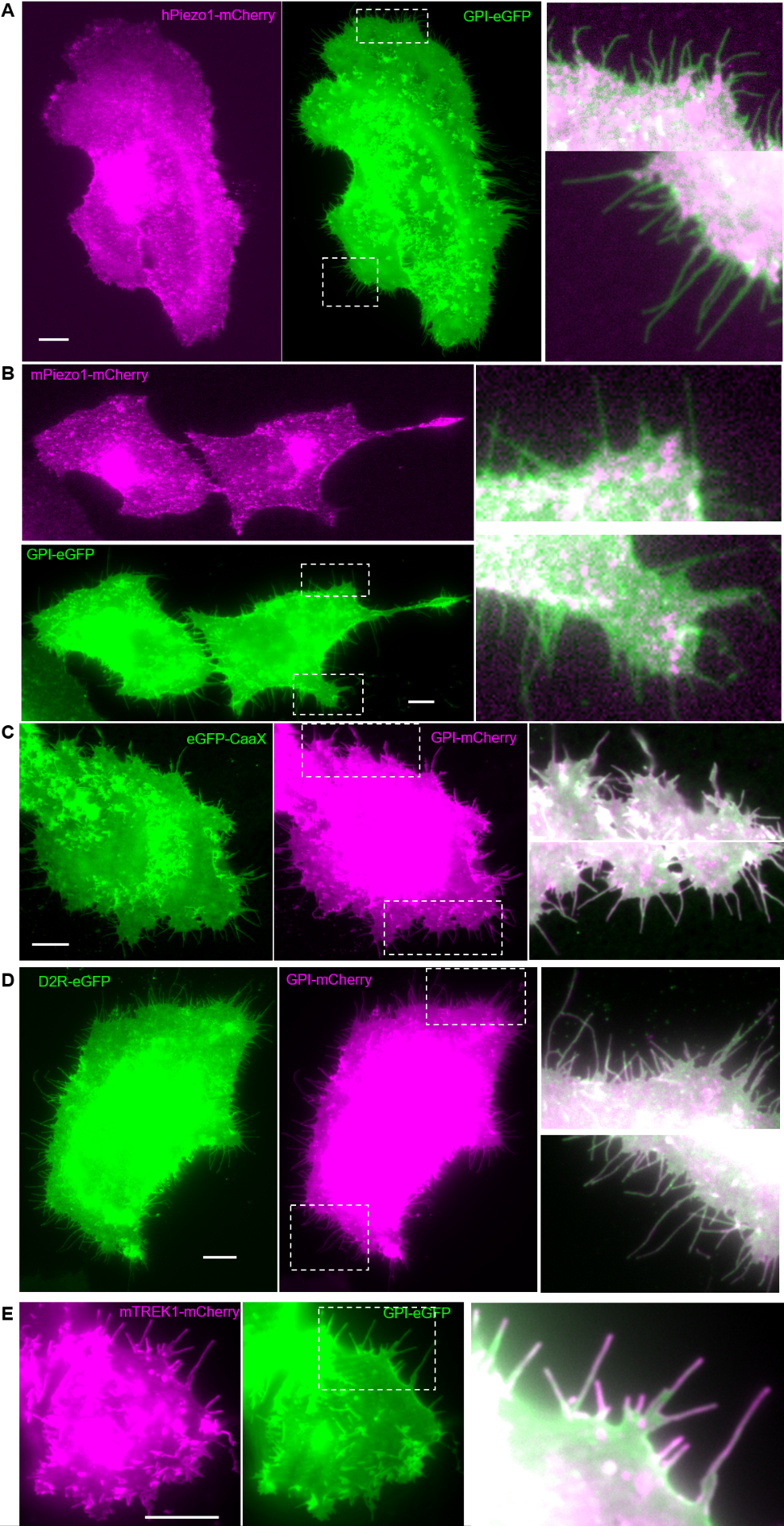
**Figure S2. Fluorescence images corresponding to Fig. 1D**

Fluorescence images of HeLa cells co-expressing: **(A)** hPiezo1-mCherry (magenta) and GPI-eGFP (green); **(B)** mPiezo1-mCherry (magenta) and GPI-eGFP (green); **(C)** eGFP-CaaX (green) and GPI-mCherry (magenta); **(D)** D2R-eGFP (green) and GPI-mCherry (magenta). **(E)** Fluorescence images of HEK-293T cells co-expressing mTREK1-mCherry (magenta) and GPI-eGFP (green). The boxed regions are merged and contrast-adjusted on the right. All scale bars are 10 µm.

**Figure S3. Calculation of membrane curvature sorting and filopodia/tether radii**

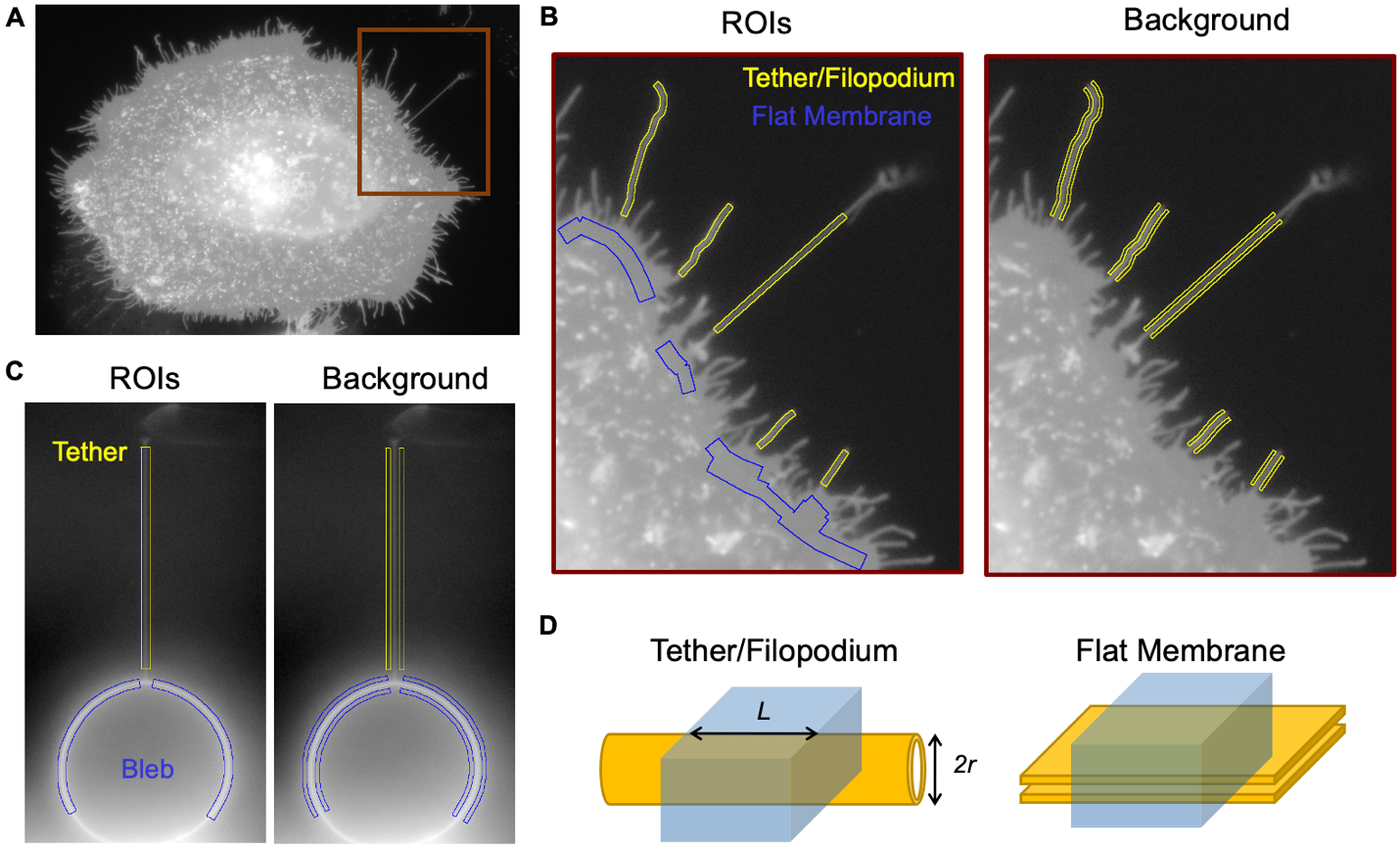

**(A)** Fluorescence images of a HeLa cell in the GPI-mCherry channel (intensity in log scale), with the boxed region enlarged in (**B**). **(B)** Left: ROIs for each tether/filopodium boxed in yellow, flat regions on the cell body boxed in blue. Right: two background regions for each ROI boxed in yellow. **(C)** Left: ROIs for a tether (yellow) and for a bleb (blue). Right: two background regions corresponding to each ROI on the left. **(D)** Illustration to show the imaged regions of a cylindrical tether/filopodium (left) and a flat cell membrane (right). Illumination profile in the ROIs shown in blue.

**Figure S4. Depletion of Piezo1 from filopodia of HEK293T cells**
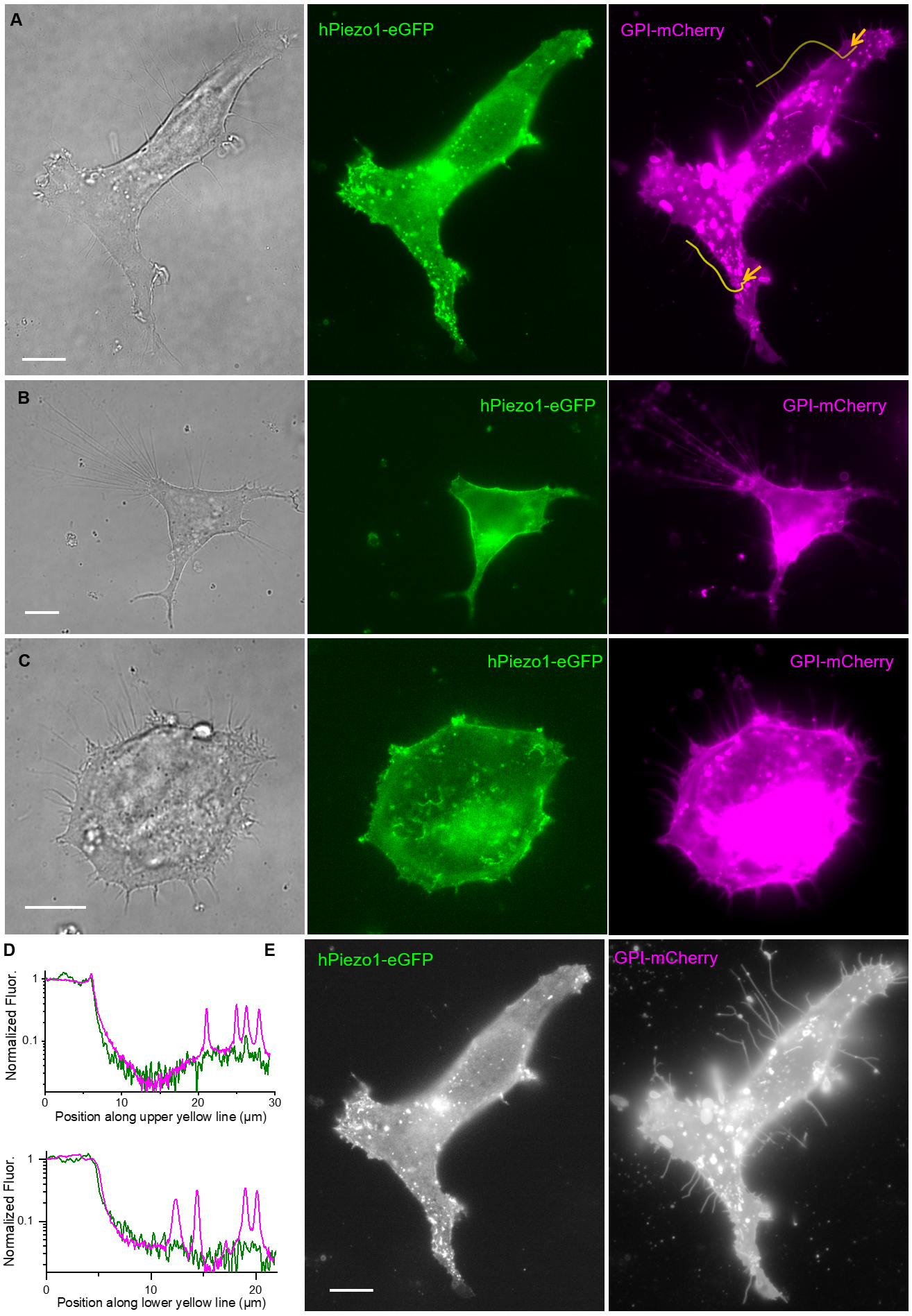

**(A-C)** Transmitted light (left), hPeizo1-eGFP fluorescence (middle), and GPI-mCherry (right) images of three representative HEK293 cells. **(D)** Fluorescent intensity along the two yellow lines shown in (**A**). **(E)** log-fluorescence of (**A**). All scale bars are 10 μm.

**Figure S5. Depletion of endogenous Piezo1 from filopodia and tether on MEFs**

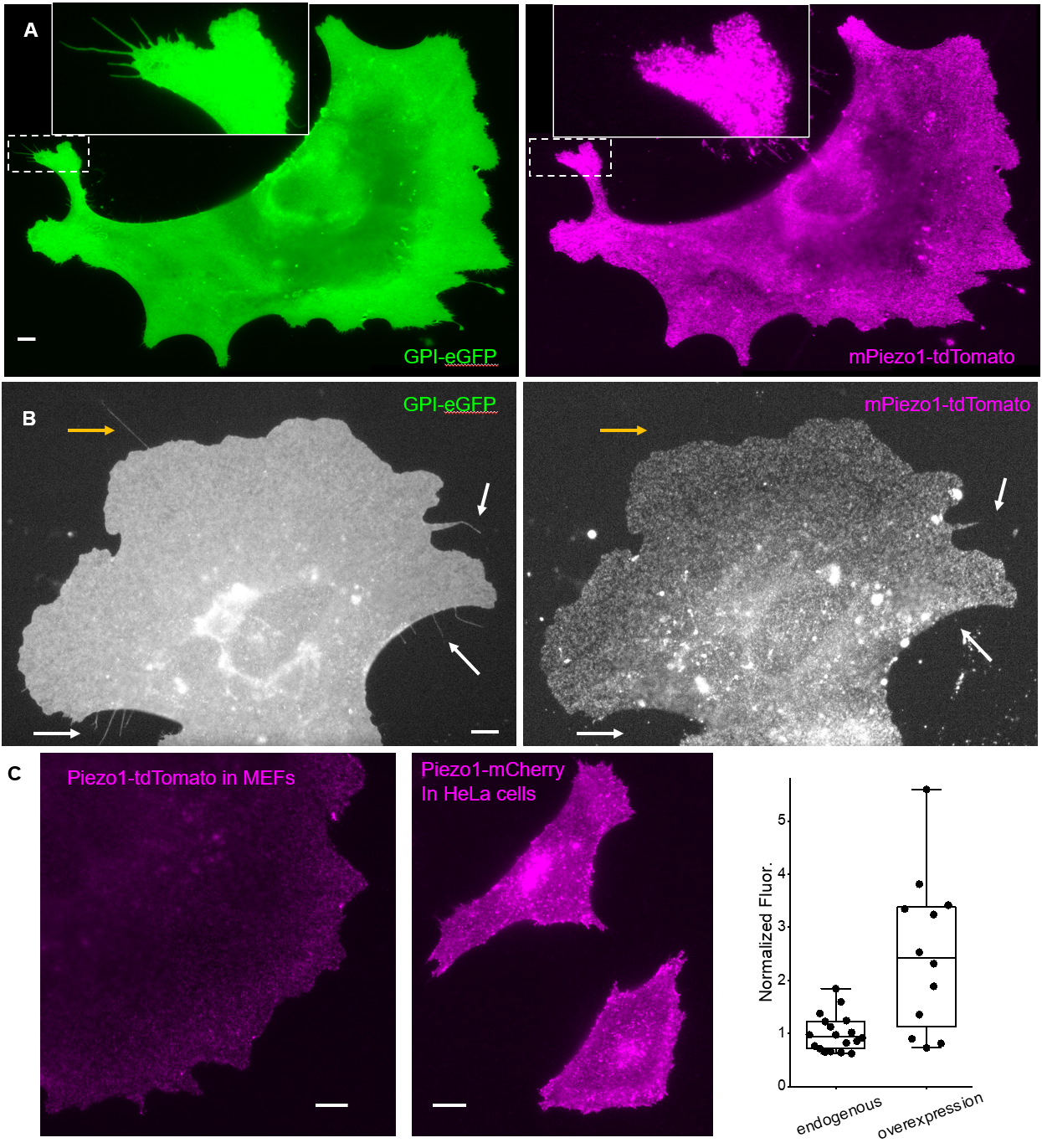
  **(A-B)** GPI-eGFP (left) and endogenous mPiezo1-tdTomato (right) fluorescence of MEFs under widefield illumination. Insets are zoom-ins of the dashed boxes. Fluorescence in (**B**) shown in log scale, white and yellow arrows in (**B**) point to filopodia and a pulled tether respectively. **(C)** Normalized fluorescence images of endogenous mPiezo1-tdTomato in MEF (left, representative of n = 18 cells) and of overexpressed hPiezo1-mCherry in HeLa (middle, representative of n = 12 cells that were chosen for Piezo sorting quantifications in Fig. 1D) under the same excitation and imaging conditions. Scattered plot of the normalized fluorescence on the right. The two images are shown with the same contrast. The normalization assumes that tdTomato is 5.94 times brighter than mCherry ^1^. All scale bars are 10 μm.

**Figure S6. Curvature sensitivity of Piezo1, D2R, and TREK1 on filopodia**

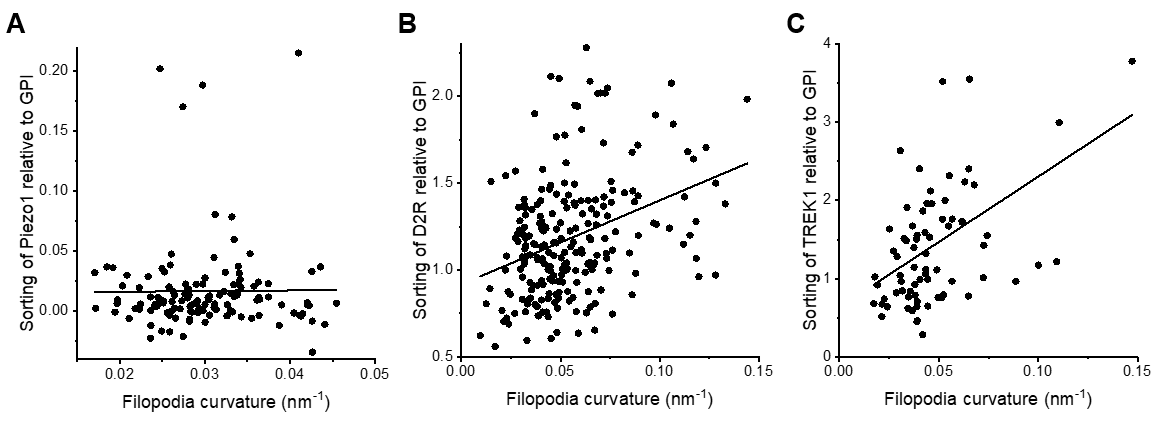

**(A)** Sorting of hPiezo1 does not change with filopodia curvature (black line: linear fit with slope 0.05 ± 0.51 nm), Person’s r value = 0.009. **(B)** Sorting of DRD2 increases with filopodia curvature (black line: linear fit with slope 4.83 ± 0.86 nm), Person’s r value = 0.35. **(C)** Sorting of TREK1 increases with filopodia curvature (black line: linear fit with slope 17 ± 4 nm), Person’s r value = 0.44.

**Figure S7. Sorting of Piezo1 does not change with the relaxation of tether radius.**

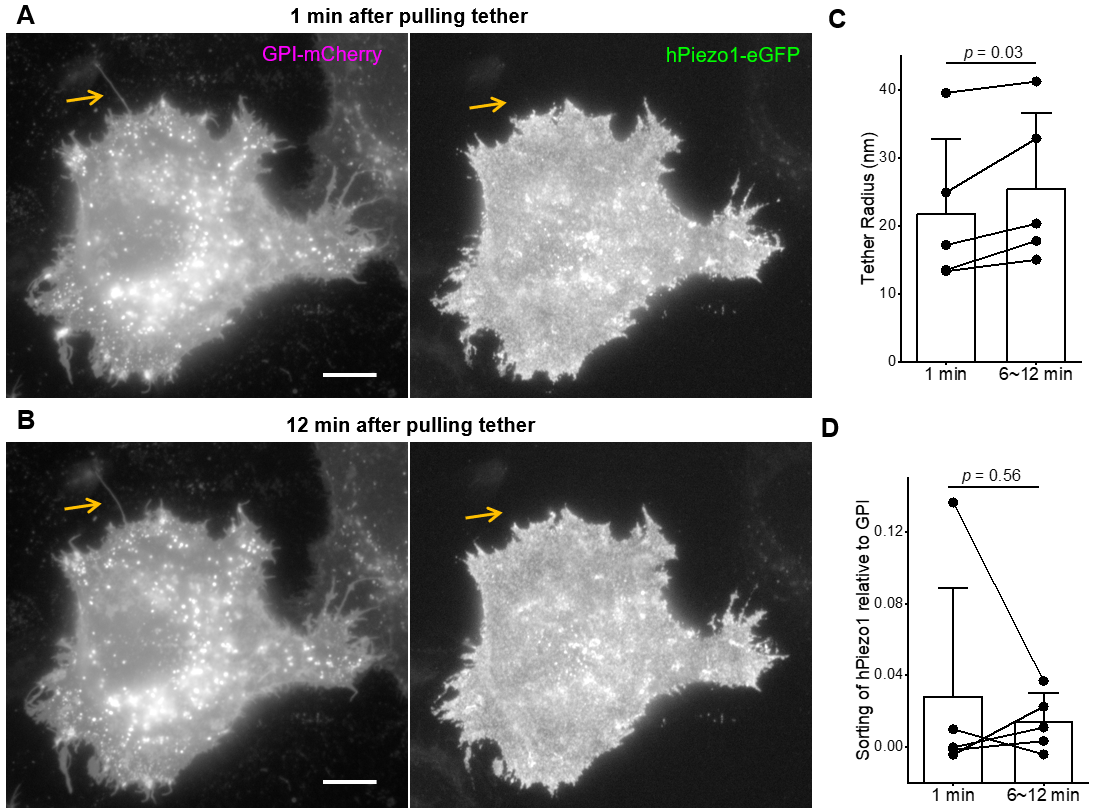

Fluorescence images of a HeLa cell co-expressing GPI-mCherry (left) and hPiezo1-eGFP (right), 1 min after pulling tether (**A**) and 12 min after pulling tether (**B**). Arrows point to the tether. The sorting of hPiezo1 was (0.010 ± 0.010) 1 min after tether pulling and was (-0.004 ± 0.011) 12 min after tether pulling, while the tether radius changed from (17.19 ± 0.05) nm to (20.33 ± 0.06) nm. Representative of 5 tethers and cells. All fluorescence images here are shown in log-scale to highlight the dim tether. All scale bars are 10 µm. Change of tether radius (**C**) and Piezo1 sorting on tethers (**D**) (1min vs. 6~12 min after tether pulling). Bar plot shows mean + SD., *p* values given by paired Student’s t test.

**Figure S8: Cell attached blebs and calculation of the upper limit for tether radii.**

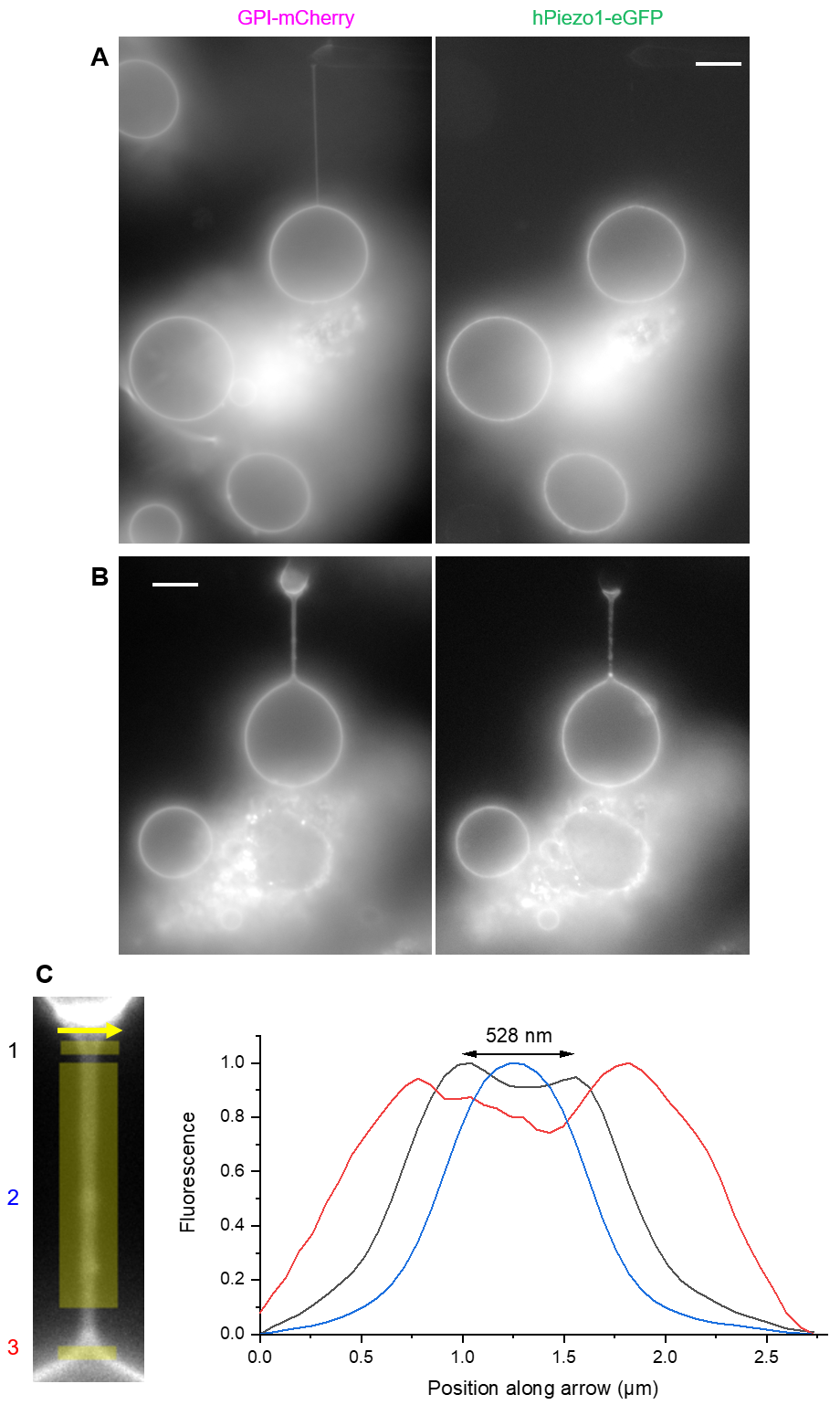
 **(A)** Image of the cell for the bleb shown in Fig. 2C. **(B)** Image of the cell for the bleb shown in Fig. 2D. All fluorescence images here are focused on the midplane of the bleb and shown in log-scale to highlight the dim tether. All scale bars are 10 µm. **(C)** Left: fluorescence image of the thickest tether pulled from a bleb (Fig. 2D). The shape of the tether gradually changes from cylindrical to catenoid-shape at the connections between the tether to the pulling handle and to the bleb, consistent with the behavior of low-tension membrane tubes ^2^. The measured apparent radius of the tether was (0.671 ± 0.003) A.U., which converts to an absolute diameter of 485 ± 82 nm. The conversion factor (361 ± 61 nm/A.U.) was determined by assuming that the average radius of filopodia and equilibrated tethers from cell body equal to the radii of tethers from tense blebs (Fig. 2G). Three yellow lines marked on the image: 1, a line across the tether-pipette junction; 2, a line across the majority of the tether (used for determining the apparent radius the tether); 3, a line across the tether-bleb junction. Right: normalized fluorescence intensity profiles along the three lines marked on the left. The majority of the tether radius was within optical resolution (blue), while a peak-to-peak distance of 528 nm was measured from the fluorescence across the tether-pipette junction (black), serving as an upper limit of the absolute diameter for this tether. A peak-to-peak distance of 1047 nm was measured from the fluorescence across the tether-bleb junction (red). The radius of the thickest tether was converted to 242 nm (Fig. 2G), consistent with the measured upper limit of tether radii (264 nm) set by line 1.

**Figure S9. Measurements of cell membrane and membrane bleb bending stiffnesses**

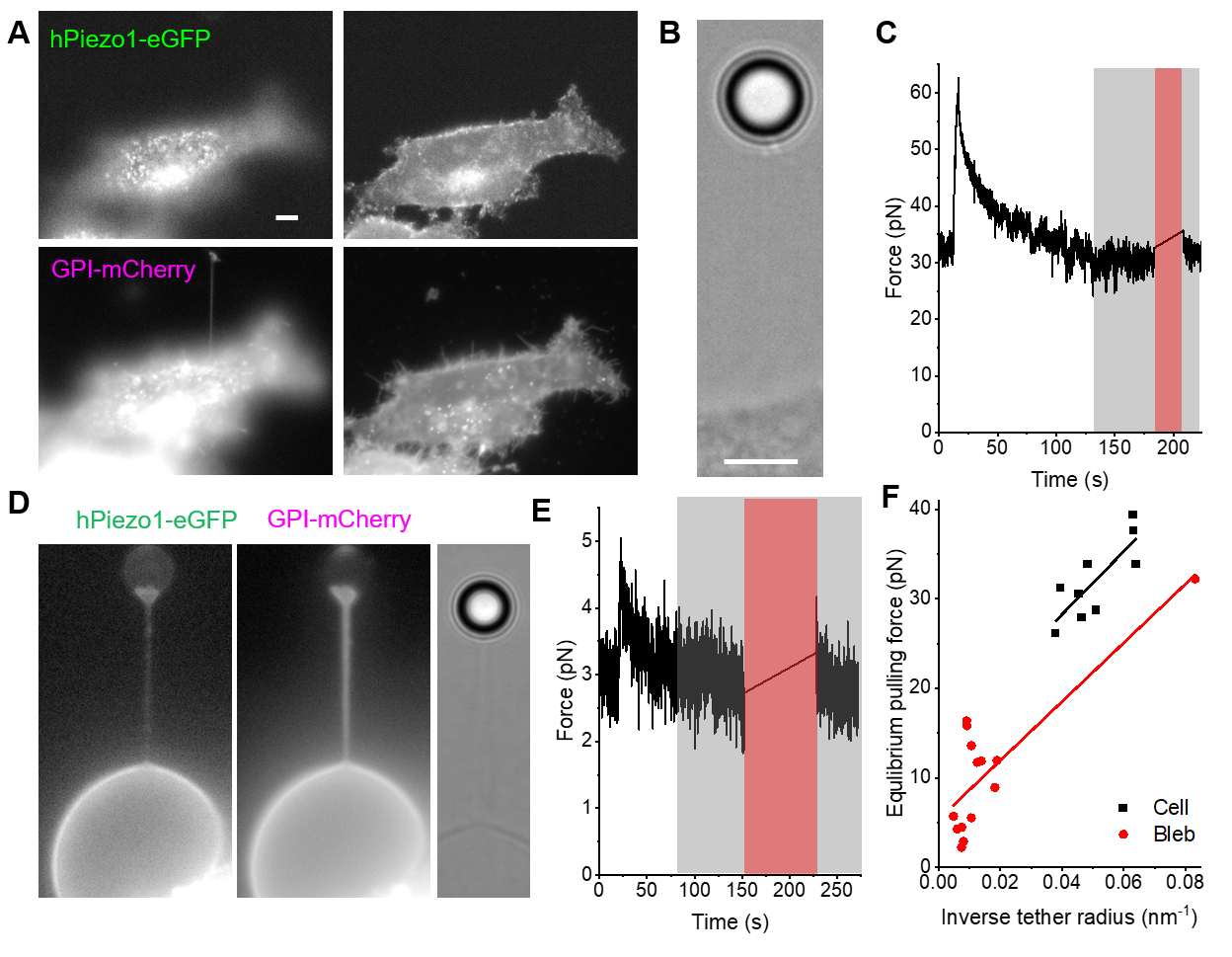

**(A)** A HeLa cell expressing hPiezo1-eGFP (up) and GPI-mCherry (down), with a 15 μm tether pulled by an optically trapped 4.5 μm diameter bead. Left: focus on the tether. Right: focus on the cell body. **(B)** Transmitted light image of the optically trapped bead that was used for calculating tether pulling force. **(C)** Time dependent tether pulling force after stretching the tether at t = 10 s. Images in (**A**) were taken during the period shaded in red, from which a 26.3 nm tether radius (based on GPI fluorescence) was determined. The gray area was used to calculate the equilibrated pulling force (30.5 pN). **(D)** A bleb from HeLa cell expressing hPiezo1-eGFP (left) and GPI-mCherry (middle), with a 15 μm tether pulled by an optically trapped 4.5 μm diameter bead (right). **(E)** Time dependent tether pulling force after stretching the tether at t = 20 s. Images in (**D**) were taken during the period shaded in red, from which a 135 nm tether radius (based on GPI fluorescence) was determined. The gray area was used to calculate the equilibrated pulling force (3 pN). **(F)** Experiments of equilibrium tether pulling force vs. inverse tether radius repeated on 9 tethers pulled from 6 independent cells (black) and 14 tethers from 14 blebs (red). Solid lines are linear fits. The slopes of the fits (black: 351 ± 99 pN·nm; red: 328 ± 63 pN·nm) gave the bending stiffness of 13.6 ± 3.8 k_B_T and 12.7± 2.5 k_B_T for cell membrane and blebs respectively. The intercepts of the fits take into account potential contributions from cytoskeletal attachments and membrane asymmetry. All scale bars are 5 μm.

**Figure S10. Modeling the curvature sorting of Piezo1**

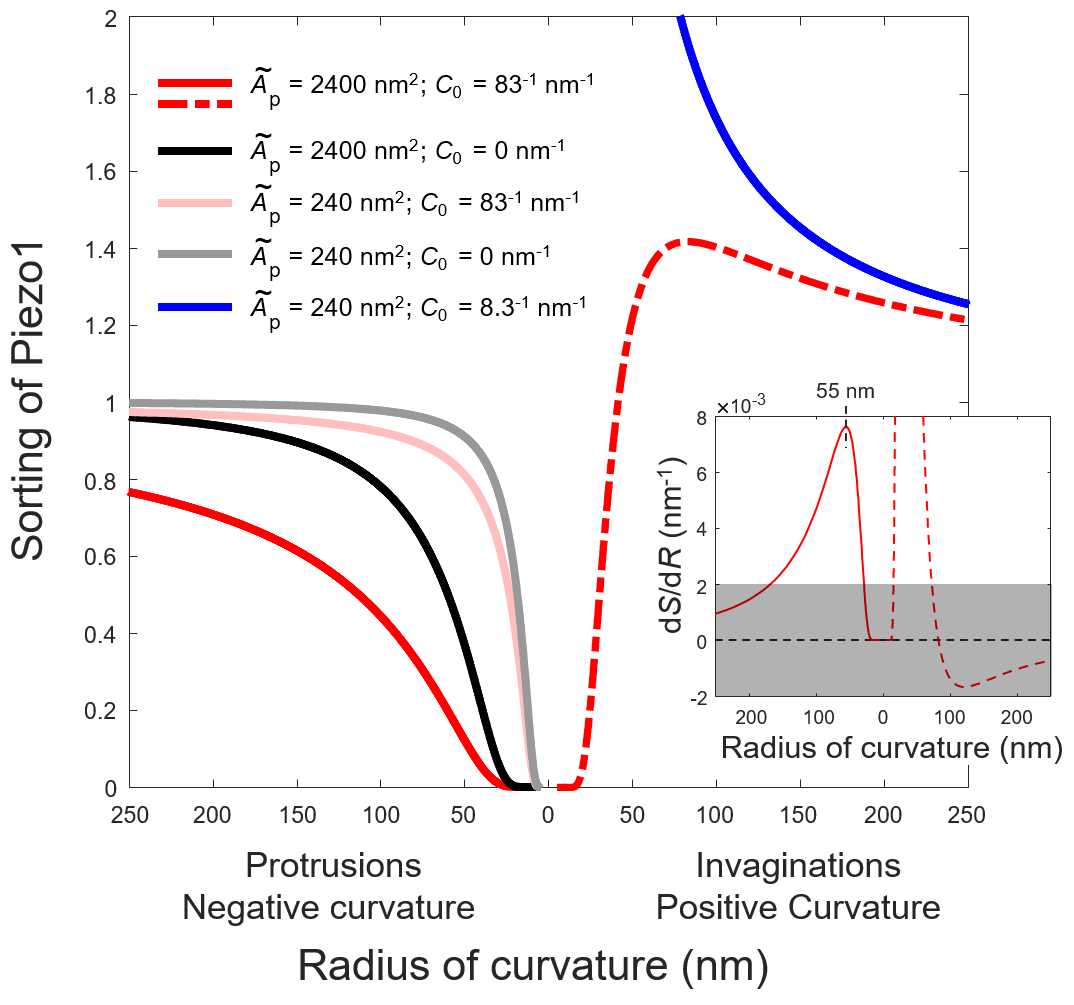

Sorting of Piezo1 as a function of membrane protrusion radius (left; eq. S14) plotted using parameters that correspond to closed Piezo1 (red, based on data in Fig. 2G) and open/inactivated Piezo1 (black, based on data in Fig. 4D). The sorting of closed Piezo1 was extrapolated to membrane invaginations (right, dashed line) according to eq. S15. The curvature sorting of a hypothetical ion channel with 1/10 of the area of Piezo1 are plotted in pink (closed) and gray (open/inactive), showing that the opening of the channel has less significant effect on its curvature sorting if the area of Piezo1 were small. The sorting of a hypothetical protein with 1/10 of the area of Piezo1 and 10 times higher spontaneous curvature is plotted in blue, showing strong curvature sensitivity to invaginations, similar to that of N-BAR domain containing proteins ^3, 4^. Inset: first derivative of the red lines, showing the sensitivity of closed Piezo1 to membrane curvature. On protrusions, the sensitivity peaks at *R*_p_ = 55 nm. The gray area indicates when the sorting of Piezo1 has low sensitivity to changes in membrane curvature (|d*S*/d*R*| < 0.002 nm^-1^).

**Figure S11. Full images for Fig. 4A, 4F**
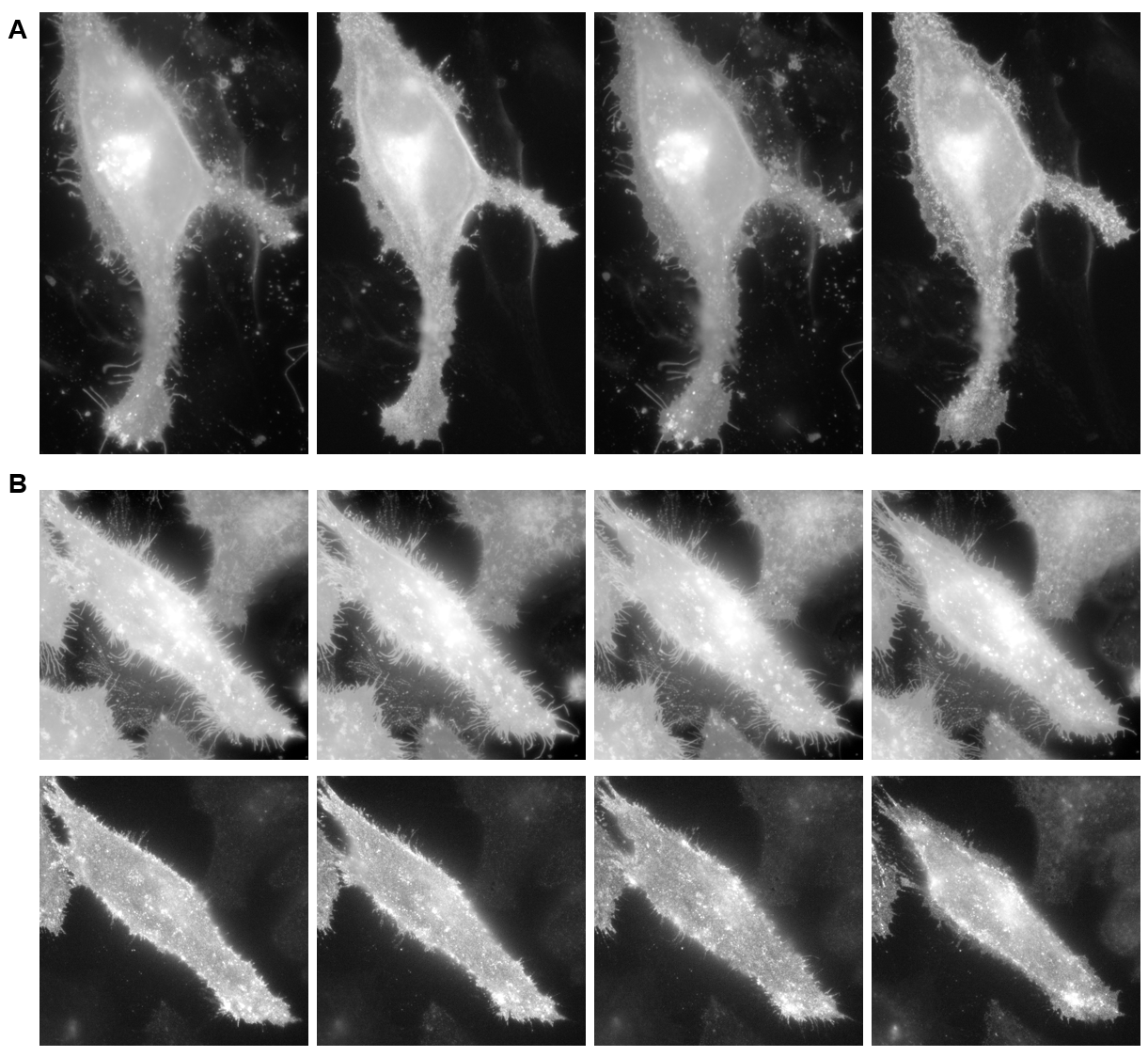

**(A)** Full image of Fig. 4A. **(B)** Full image of Fig. 4F

**Figure S12. Kinetics of hPiezo1 on plasma membranes**

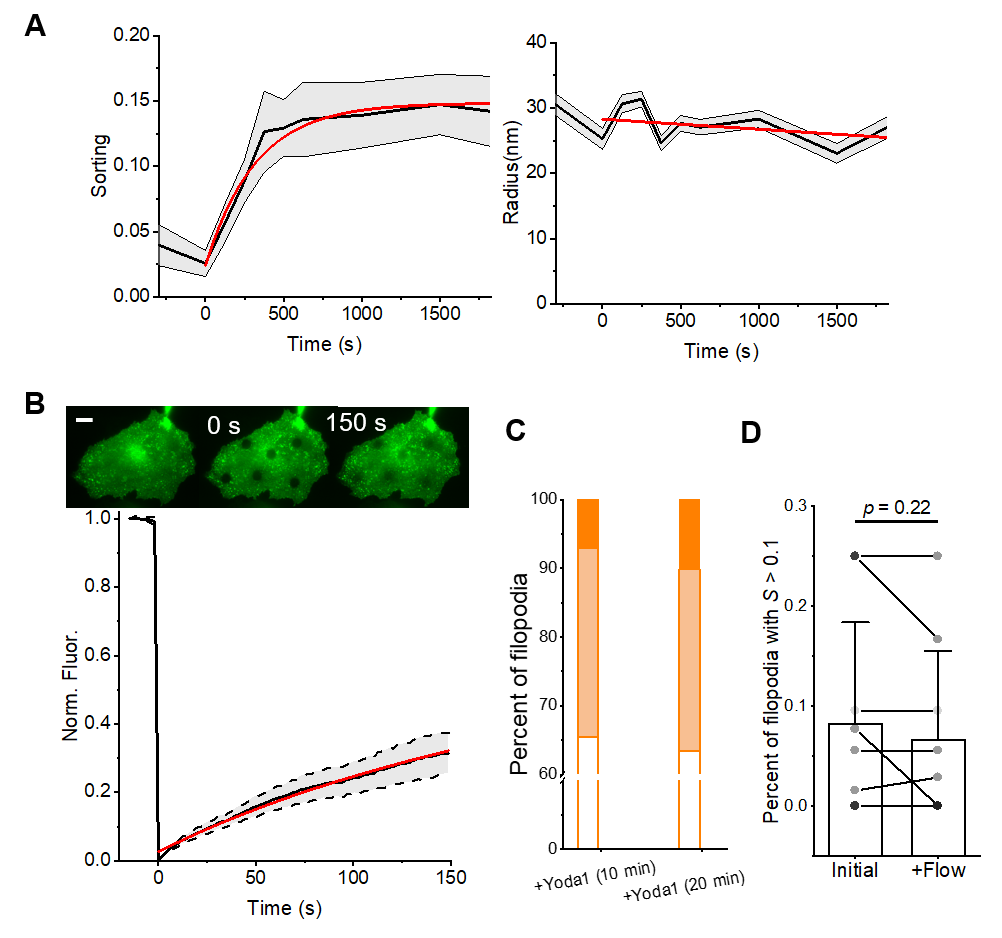

(**A**) Change of Piezo1 sorting on filopodia (left) and filopodia radii (right) after adding 100 µM Yoda1. Error bars are standard error of the mean. A time constant of 300 ± 40 s was obtained by a single exponential fit to the sorting data (red; left). A slope of -0.0015 ± 0.0015 nm/s was obtained by fitting the radius data to a line (red; right). (**B**) Fluorescence recovery after photobleaching (FRAP) of hPiezo1-eGFP in HeLa cells. Red line: fitting to eq. S8, $\tau_{0.5}$ = 342 ± 4 s, R^2^ = 0.994. Error bars are standard deviation. Scale bar, 5 μm. (**C**) Percentage of filopodia that showed strong (S_filo_ > 0.3, dark), weak (0.1 < S_filo_ < 0.3, light), and no (S_filo_ < 0.1, open) sorting of hPiezo1 10 min and 20 min after adding Yoda1. 10 µM Yoda1 was added after hypotonic shock. **(D)** Fraction of filopodia with measurable sorting of Piezo1 tracked on 9 cells under shear stress (applied by XC medium only) that mimics the strongest washing steps in Fig. 4E-4G. Initial: before applying flow. +Flow: 10 min after applying flow. *p* value given by paired Student’s t-test.

stylefix
